# Development of a Comprehensive Analytical Workflow for eDNA Analysis of Vertebrates, Arthropods, and Mollusks

**DOI:** 10.1101/2025.04.09.643821

**Authors:** Mio Omino, Ibuki Sugawara, Kentaro Takada, Kazutoshi Yoshitake

## Abstract

Environmental DNA (eDNA) analysis is a powerful tool for biodiversity monitoring, but conventional methods are often limited to specific taxonomic groups and short-read sequencing. This study aimed to enhance species detection using Oxford Nanopore long-read sequencing and 16Sar/br primers. Our results showed that MiFish primers effectively detected fish species, while COI primers primarily detected bacterial DNA, limiting eukaryotic identification. In contrast, 16Sar/br primers enabled the detection of diverse taxa, including fish, mollusks, and crustaceans.

To improve accuracy, we expanded the mitochondrial 16S database, increasing its size from 4.7 GB to 7.0 GB and the number of sequences from 1.69M to 2.15M, leading to improved mollusk detection. Additionally, an automated analysis pipeline was developed, streamlining data processing. These findings highlight the potential of long-read sequencing and database enhancement for more comprehensive eDNA-based biodiversity assessments.

## Introduction

In recent years, the application of environmental DNA (eDNA) analysis has rapidly increased in ecology and conservation. The presence/absence data obtained from eDNA allows large-scale monitoring of populations and identification of habitats valuable for species of conservation concern. For example, Ficetola et al. (2008) successfully detected bullfrogs using eDNA, leading to numerous subsequent studies^1^. Moreover, compared to traditional survey methods, eDNA analysis offers benefits such as non-invasive monitoring and reduced survey effort^2^.

Research utilizing environmental DNA metabarcoding for freshwater fish has demonstrated the utility and limitations of sample pooling and has also shown that terrestrial mammals can be detected from the water of forest ponds. With further generalization, eDNA analysis is expected to be applied to fisheries resource data collection and invasive species monitoring. Currently, eDNA analysis primarily targets specific taxonomic groups, such as fish, birds, and mammals^3, 4^. However, universal primers capable of detecting multiple taxa in a single sequencing run have not been well established. Moreover, conventional eDNA analysis mainly employs short-read sequencing using the next-generation sequencer Illumina MiSeq, which limits sequence length^4^. Given these limitations, this study aims to establish a more comprehensive species detection method by amplifying the 16S gene region using the 16S ar, br primers, traditionally used for species identification with Sanger sequencing, and conducting long-read sequencing with Oxford Nanopore.

## Results

First, we performed PCR with MiFish primers to determine the detectable species. The results showed that fish species expected to inhabit the sampling sites were comprehensively detected (Figure 1, Table 1). Next, PCR amplification was conducted using LCO1490 and HCO2198 primers, targeting the COI region, to confirm detectable species. However, most of the detected sequences were assigned to bacteria (Figure 2), indicating that COI-based eDNA analysis cannot specifically detect eukaryotic DNA in environments with abundant bacteria, such as seawater.

**Figure 1.**
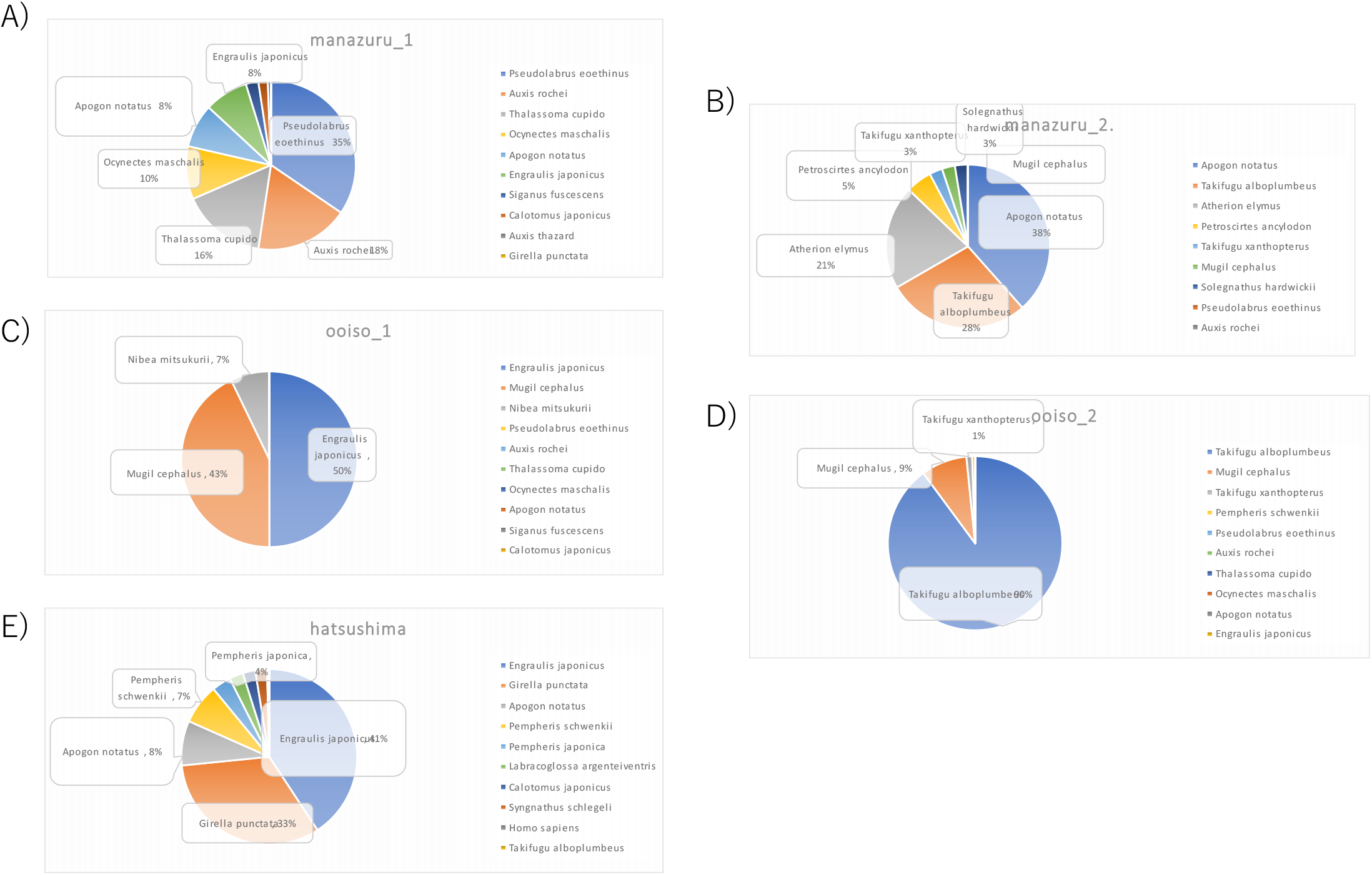
The result of eDNA analysis of the MiFish primer Water samples were collected from the coasts of Manazuru (A, B), Oiso (C, D) and Hatsushima (E). eDNA analysis was conducted using the MiFish primer.

**Figure 2.**
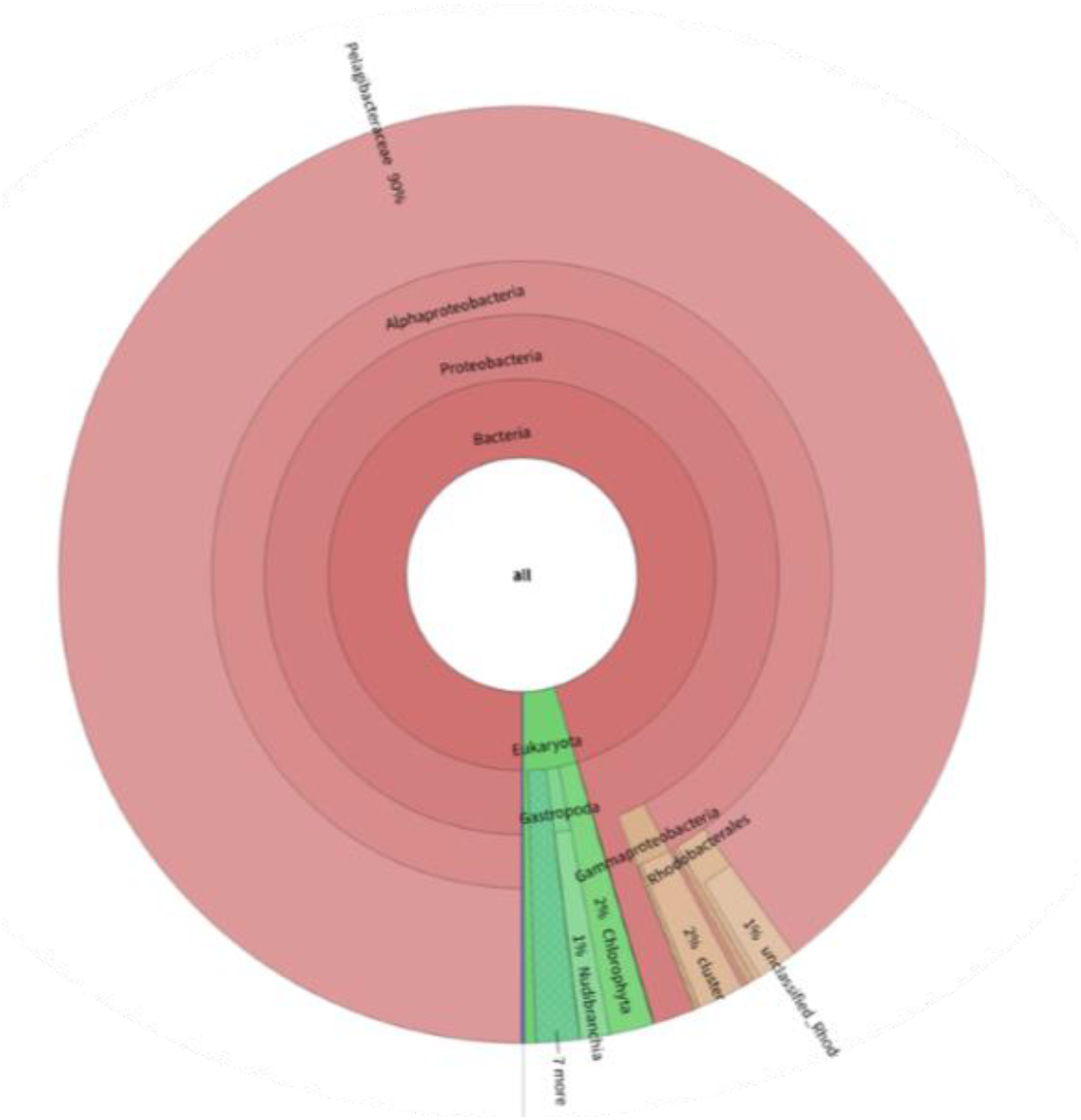
The result of eDNA analysis of the COI primer Water samples were collected from the coasts of Atami.

**Table 1.** The result of eDNA analysis of the MiFish primer.

| #Datasets | ogasawara | manazuru | manazuru | ooiso_1 | ooiso_2 | hatsushin |
| --- | --- | --- | --- | --- | --- | --- |
|  | 100 | 100 | 100 | 100 | 100 | 100 |
| *Pseudolabrus eoethinus* | 0 | 34.52381 | 0 | 0 | 0 | 0 |
| *Auxis rochei* | 0 | 17.85714 | 0 | 0 | 0 | 0 |
| *Thalassoma cupido* | 0 | 16.07143 | 0 | 0 | 0 | 0 |
| *Ocynectes maschalis* | 0 | 10.11905 | 0 | 0 | 0 | 0 |
| *Apogon notatus* | 0 | 8.333333 | 38.46154 | 0 | 0 | 8.232119 |
| *Engraulis japonicus* | 0 | 8.333333 | 0 | 50 | 0 | 40.75574 |
| *Siganus fuscescens* | 0 | 2.380952 | 0 | 0 | 0 | 0 |
| *Calotomus japonicus* | 0 | 1.785714 | 0 | 0 | 0 | 2.294197 |
| *Auxis thazard* | 0 | 0.595238 | 0 | 0 | 0 | 0 |
| *Girella punctata* | 0 | 0 | 0 | 0 | 0 | 32.65857 |
| *Labracoglossa argenteiventris* | 0 | 0 | 0 | 0 | 0 | 2.564103 |
| *Nibea mitsukurii* | 0 | 0 | 0 | 7.142857 | 0 | 0 |
| *Pempheris japonica* | 1.515152 | 0 | 0 | 0 | 0 | 3.508772 |
| *Pempheris schwenkii* | 1.515152 | 0 | 0 | 0 | 0.531915 | 7.422402 |
| *Takifugu alboplumbeus* | 71.21212 | 0 | 28.20513 | 0 | 89.89362 | 0.134953 |
| *Takifugu xanthopterus* | 0 | 0 | 2.564103 | 0 | 1.06383 | 0 |
| *Atherion elymus* | 0 | 0 | 20.51282 | 0 | 0 | 0 |
| *Petroscirtes ancylodon* | 0 | 0 | 5.128205 | 0 | 0 | 0 |
| *Mugil cephalus* | 0 | 0 | 2.564103 | 42.85714 | 8.510638 | 0 |
| *Solegnathus hardwickii* | 0 | 0 | 2.564103 | 0 | 0 | 0 |
| *Syngnathus schlegeli* | 0 | 0 | 0 | 0 | 0 | 2.159244 |
| *Homo sapiens* | 25.75758 | 0 | 0 | 0 | 0 | 0.269906 |

Subsequently, PCR amplification targeting the 16S gene region using 16Sar/br primers successfully detected a wide range of species, including fish, mollusks, and crustaceans, expected to inhabit the sampling sites (Figure 3). However, species match rates were approximately 85%, indicating low reliability, likely due to the absence of numerous species in the initial database.

**Figure 3.**
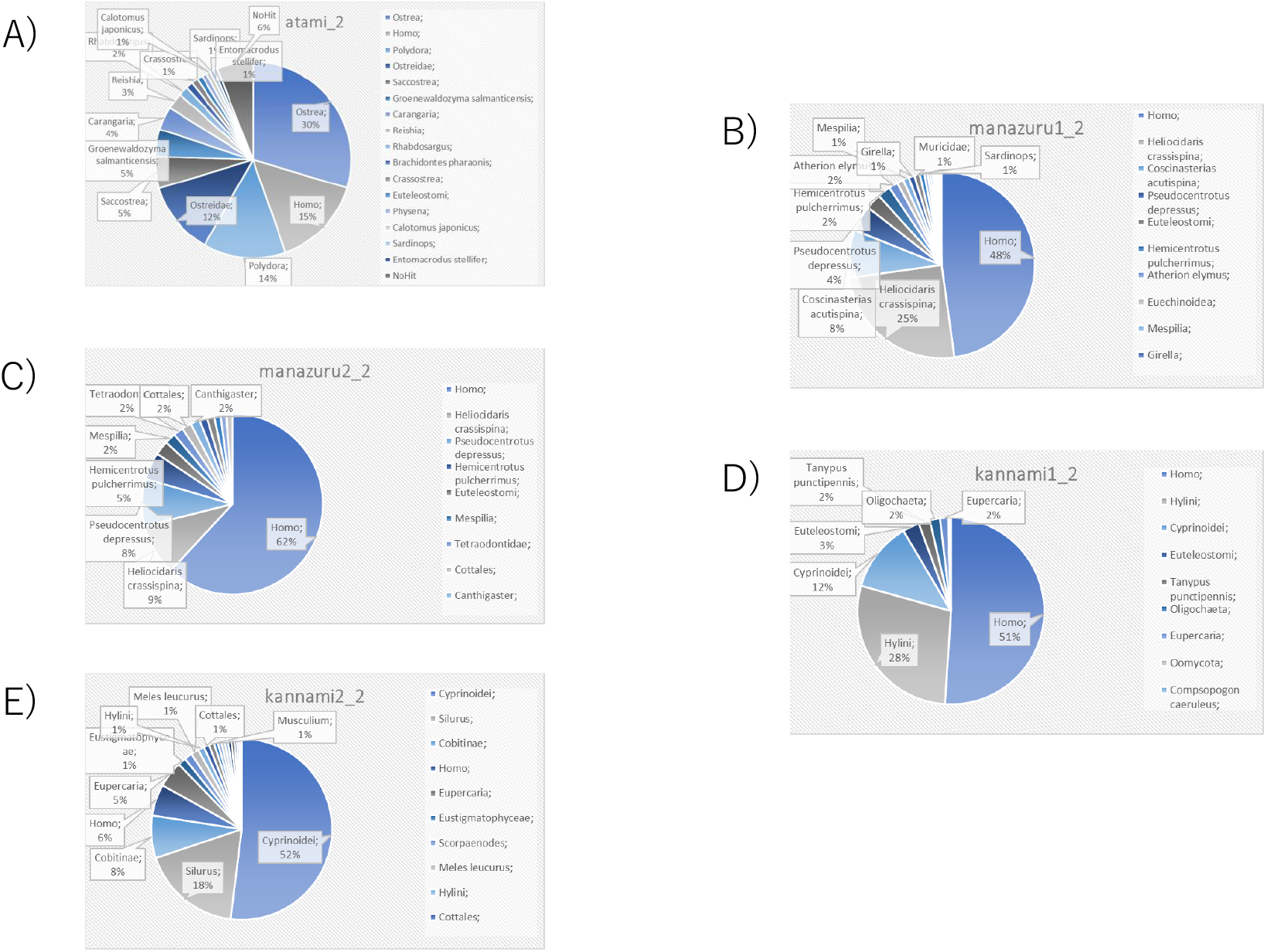
The result of eDNA analysis of the 16Sar/br primer Water samples were collected from the coasts of Atami (A), Manazuru (B, C) and Kannami (D, E). eDNA analysis was conducted using the 16Sar/br primer.

The BLAST search database used included MitoFish (12S rRNA), Silva (bacterial 16S rRNA and eukaryotic 18S rRNA), and NCBI mitochondrial full-length sequences, but it lacked partial sequences of the mitochondrial 16S rRNA. To improve the accuracy and increase the number of detected species, we added mitochondrial 16S rRNA data to the BLAST search database. As a result, the database size increased from 4.7 GB to 7.0 GB, and the number of entries increased from 1,698,849 to 2,151,945. A comparative BLAST search revealed that sequences previously unidentifiable at a 95% similarity threshold using the existing database were now classified as mollusks, primarily bivalves (Table 2, 3).

**Table 2.** Comparison of the analysis of the Kannami sample using the old and new databases.

|  | new database | old database |
| --- | --- | --- |
| Total hits | 4119 | 3611 |
| *Physarum polycephalum* | 1 | 0 |
| *Silurus asotus* | 11 | 7 |
| *Misgurnus anguillicaudatus* | 216 | 207 |
| *Gnathopogon elongatus* | 49 | 49 |
| *Nipponocypris koreanus* | 0 | 1 |
| *Nipponocypris sieboldii* | 1 | 0 |
| *Nipponocypris temminckii* | 2623 | 2564 |
| *Culter oxycephaloides* | 1 | 1 |
| *Homo heidelbergensis* | 0 | 2 |
| *Homo sapiens* | 1042 | 670 |
| *Meles anakuma* | 104 | 0 |
| *Meles leucurus* | 0 | 96 |
| *Dryophytes japonicus* | 47 | 14 |
| *Ephemera japonica* | 1 | 0 |
| *Ephemera* sp. EP186 | 13 | 0 |
| *Dendostrea* sp. 4 CL-2021 | 1 | 0 |
| *Semisulcospira dolorosa* | 1 | 0 |
| *Semisulcospira libertina* | 7 | 0 |
| *Semisulcospira trachea* | 1 | 0 |

**Table 3.**
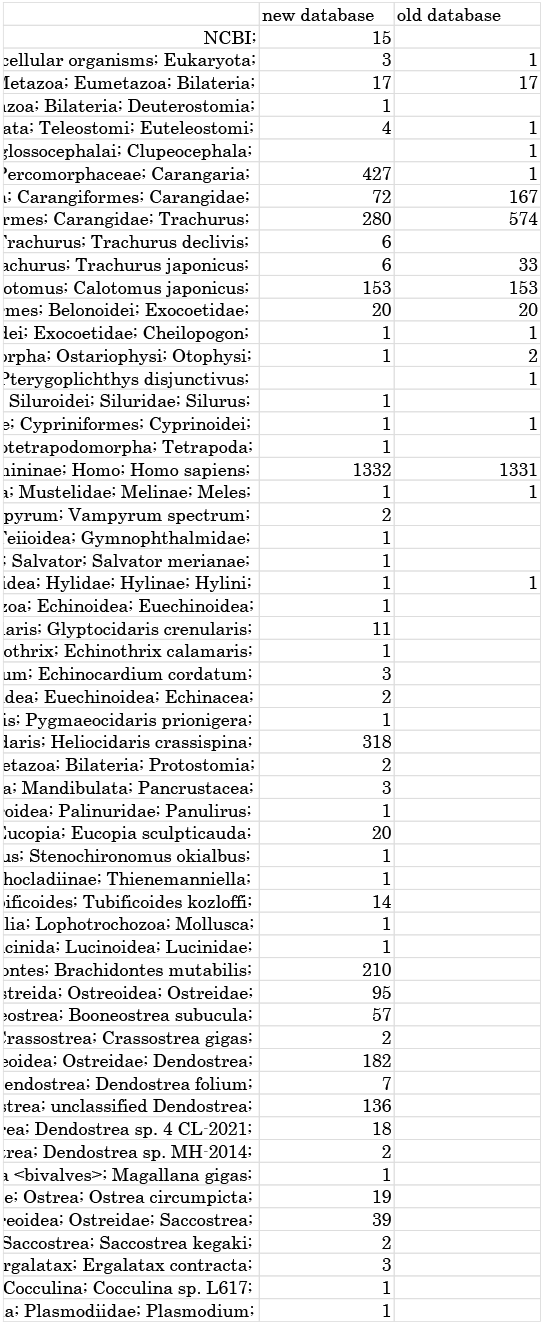
Comparison of the analysis of the Atami sample using the old and new databases.

While the MiFish primer could detect only about 10 fish species (Table 1), the 16Sar/br universal primers enabled the detection of 16–54 species, including mollusks, amphibians, and mammals, in a single sequencing run (Table 2, 3). Additionally, using 16Sar/br primers, 18 out of 24 fish species detected with MiFish primers were also identified.

## Methods

### Nanopore Sequencing

Seawater samples (2 L) were filtered through a 0.45 _μ_m membrane filter, and DNA was extracted following the eDNA Society Manual. Each sample was treated with 400 μL of Buffer AL and 40 μL of Proteinase-K, incubated at 56°C for 30 minutes, and centrifuged at 500g for 20 seconds. DNA was purified using the DNeasy Blood and Tissue Kit, and DNA concentration was measured using NanoDrop Lite.

PCR was performed in a 10 μL reaction with 1 μL sterile distilled water, 1.5 μL of each primer (0.3 μM) (Table 4), 5 μL of repliQa HiFi ToughMix, and 1 μL of template DNA, for 45 cycles. The thermocycling conditions were: initial denaturation at 95°C for 3 minutes, denaturation (98°C, 10 sec), annealing (55°C, 5 sec), and extension (68°C, 1 sec) for 45 cycles and final extension at 68°C for 1 minute.

**Table 4.**
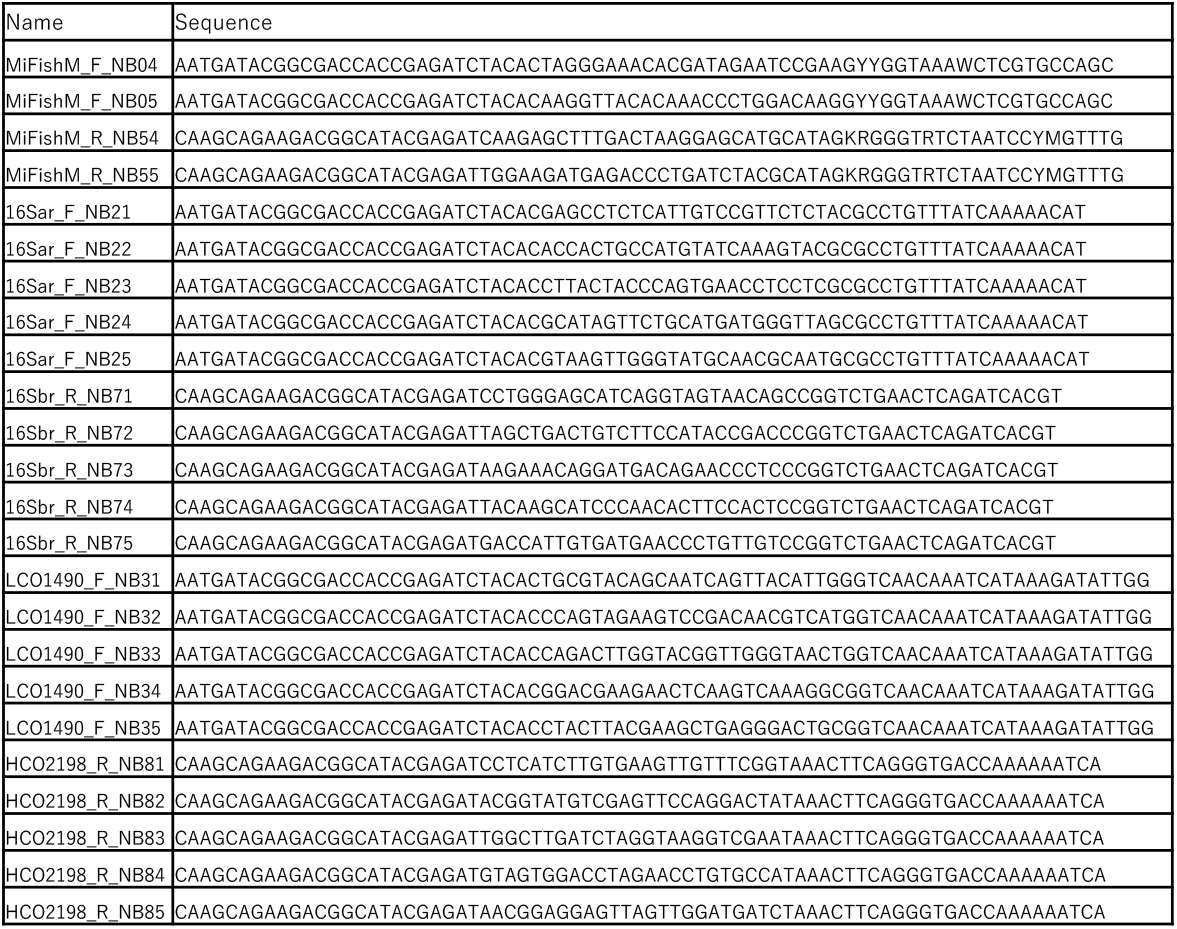
Primer list.

PCR products were electrophoresed using 2% agarose gel, and target bands were extracted and purified using FastGene Gel/PCR Extraction Kit (Qiagen). Libraries were prepared according to Oxford Nanopore Technologies’ protocol, and sequencing was performed on a Flongle R10.4.1 flow cell.

#### Database Construction

Representative mitochondrial sequences were obtained from the literature^5-13^. Sequences were categorized based on whether the 16S gene region was identified. Accession IDs and start/end positions were used to extract only the 16S gene region, forming FASTA files. For sequences without specific 16S gene annotations, BioPython was used to identify the 16S region from GenBank, and corresponding FASTA files were created.

A database was generated by removing duplicates and retrieving FASTA sequences from the NCBI nt database (https://github.com/mio49204920/eDNA).

### Analytical Workflow

Oxford Nanopore sequencing offers long-read capabilities, real-time analysis, and cost efficiency, contributing to the generalization of eDNA analysis. However, sequencing analysis involves multiple steps. To streamline the process, an automated shell script was developed (https://github.com/mio49204920/eDNA).

The workflow includes: conversion FASTQ to FASTA files, BLAST similarity search, phylogenetic analysis using MEGAN and result compilation.

## Discussion

eDNA analysis is increasingly utilized for biodiversity conservation and resource management. However, species detectability is limited, and analytical environments remain insufficient. This study focused on mitochondrial 16S rRNA to comprehensively detect eukaryotic DNA in seawater and conducted database reinforcement and automation.

Results indicated that relevant species were detected with the 16S rRNA but bacteria were primarily detected with the COI gene region, demonstrating its inability to specifically detect eukaryotes in seawater. If COI-based eDNA analysis can be refined, it may enhance mollusk detection. Future work will focus on developing COI primers specific to eukaryotic DNA.

Database enhancement increased species detection at >95% identity, enabling high-precision identification of previously undetectable mollusks and crustaceans. Further database refinement will be conducted to eliminate redundant data and improve accuracy. Automation significantly reduced analysis time and simplified workflows, making eDNA analysis accessible to a broader audience and contributing to its widespread adoption.

## Data availability

All sequencing data are available in the Sequence Read Archive (SRA) of NCBI under the accession numbers PRJNA1229139 and PRJNA1232963.

## References

1. Gentile Francesco Ficetola, Claude Miaud, Francoise Pompanon, Pierre Taberlet. Species detection using environmental DNA from water samples. biology Letters, 2008; 423–425

2. Helen C. Rees, Ben C. Maddison, David J. Middleditch, James R.M. Patmore, Kevin C. Gough. The detection of aquatic animal species using environmental DNA – a review of eDNA as a survey tool in ecology. Journal of Applied Ecology, Vol. 51, 2014; 1450–1459

3. M. Miya, Y. Sato, T. Fukunaga, T. Sado, J. Y. Poulsen, K. Sato, T. Minamoto, S. Yamamoto, H. Yamanaka, H. Araki, M. Kondoh and W. Iwasaki. MiFish, a set of universal PCR primers for metabarcoding environmental DNA from fishes: detection of more than 230 subtropical marine species. Royal Society Open Science, Vol. 2, 2015; 1–16

4. Masayuki Ushio, Koichi Murata, Tetsuya Sado, Isao Nishiumi, Masamichi Takeshita, Wataru Iwasaki & Masaki Miya. Demonstration of the potential of environmental DNA as a tool for the detection of avian species. Scientific Reports 8, 2018

5. Li Yang, Zongqing Tan, Daren Wang, Ling Xue, Min-xin Guan, Taosheng Huang & Ronghua Li. Species identification through mitochondrial rRNA genetic analysis. Scientific Reports 4, 2014

6. Takashi P. Satoh, Masaki Miya, Kohji Mabuchi & Mutsumi Nishida. Structure and variation of the mitochondrial genome of fishes. BMC Genomics 17, 2016

7. Bingjian Liu, Ying Peng, Kun Zhang, Yifan Liu, Jiasheng Li, Jian Chen, Xuepeng Li, Xun Jin, Sixu Zheng and Yunpeng Wang. Comparative analysis of the complete mitochondrial genomes of two species of Clupeiformes and the phylogenetic implications for Clupeiformes. Cambyidge Core, Vol. 102, 2022

8. Wenjing Li, Ning Qiu, Hejun Du. Complete mitochondrial genome of Rhodeus cyanorostris (Teleostei, Cyprinidae): characterization and phylogenetic analysis. ZooKeys, 2022; 111–125

9. Hoyoung Chung. Phylogenetic analysis and characterization of mitochondrial DNA for Korean native cattle. Open Journal of Genetics, Vol. 3, 2013

10. Hu Li Yan, Juan Li. Eighteen mitochondrial genomes of Syrphidae (Insecta: Diptera: Brachycera) with a phylogenetic analysis of Muscomorpha. PLOS one, 2023

11. Rui-Wen Wu, Xiong-Jun Liu, Sa Wang, Kevin J. Roe, Shan Ouyang, Xiao-Ping Wu. Analysis of mitochondrial genomes resolves the phylogenetic position of Chinese freshwater mussels (Bivalvia, Unionidae). ZooKeys, 2019; 23–46

12. Kin Onn Chan, Stefan T. Hertwig, Dario N. Neokleous, Jana M. Flury & Rafe M. Brown. Widely used, short 16S rRNA mitochondrial gene fragments yield poor and erratic results in phylogenetic estimation and species delimitation of amphibians. BMC Ecology and Evolution 22, 2022

13. Federico Plazzi, Marco Passamonti. Towards a molecular phylogeny of Mollusks: Bivalves’ early evolution as revealed by mitochondrial genes. Molecular Phylogenetics and Evolution, Vol. 57, 2010; 641-657

